## Supplementary material for "Noncanonical usage of stop codons in ciliates expands proteins with Q-rich motifs": This file has three figures and five tables

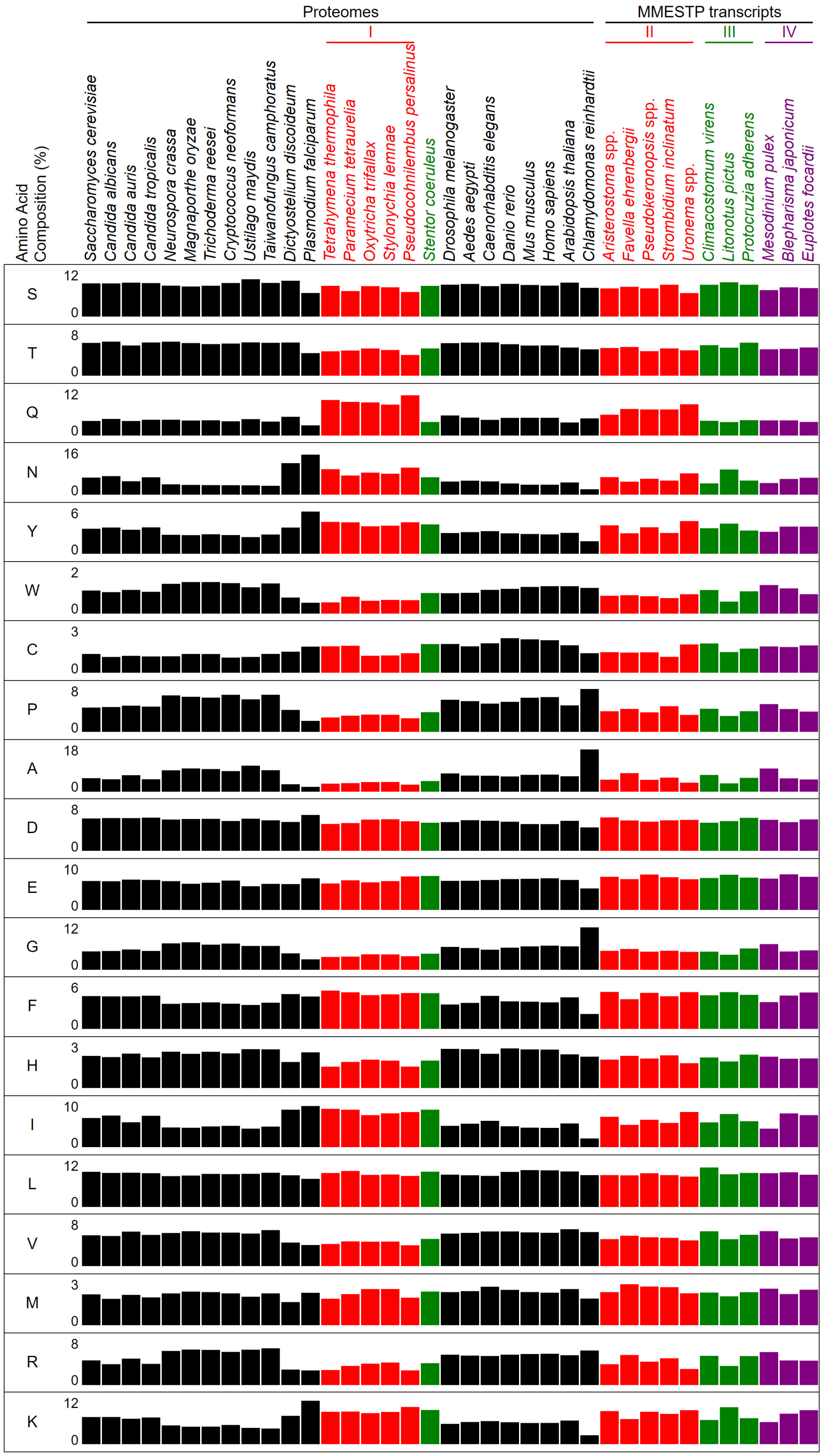


**Figure S1. Proteome-wide contents of 20 different amino acids in 37 different eukaryotes.**

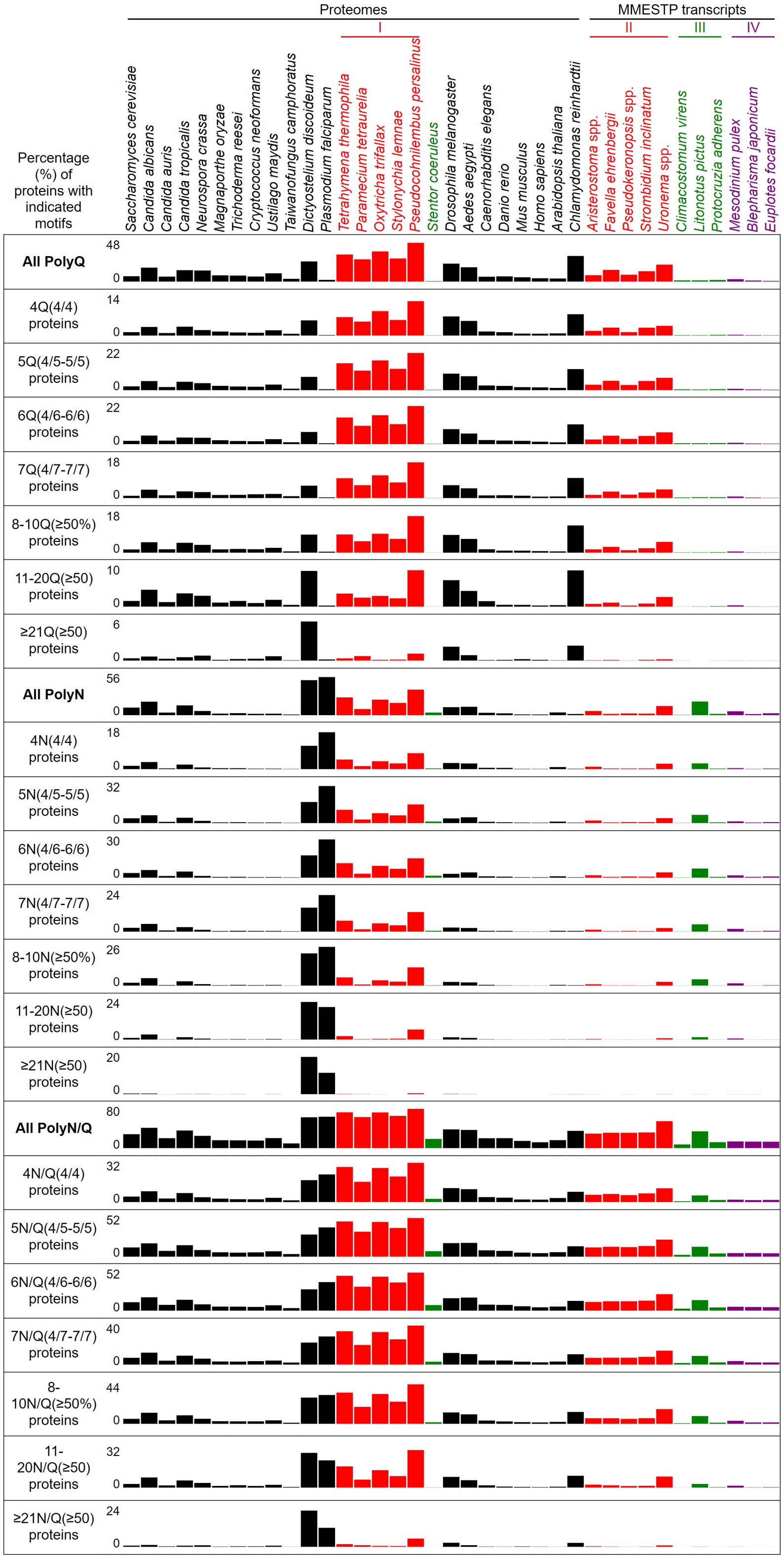


**Figure S2. Percentages of proteins with indicated polyQ and polyQ/N tracts in 37 different eukaryotes.**

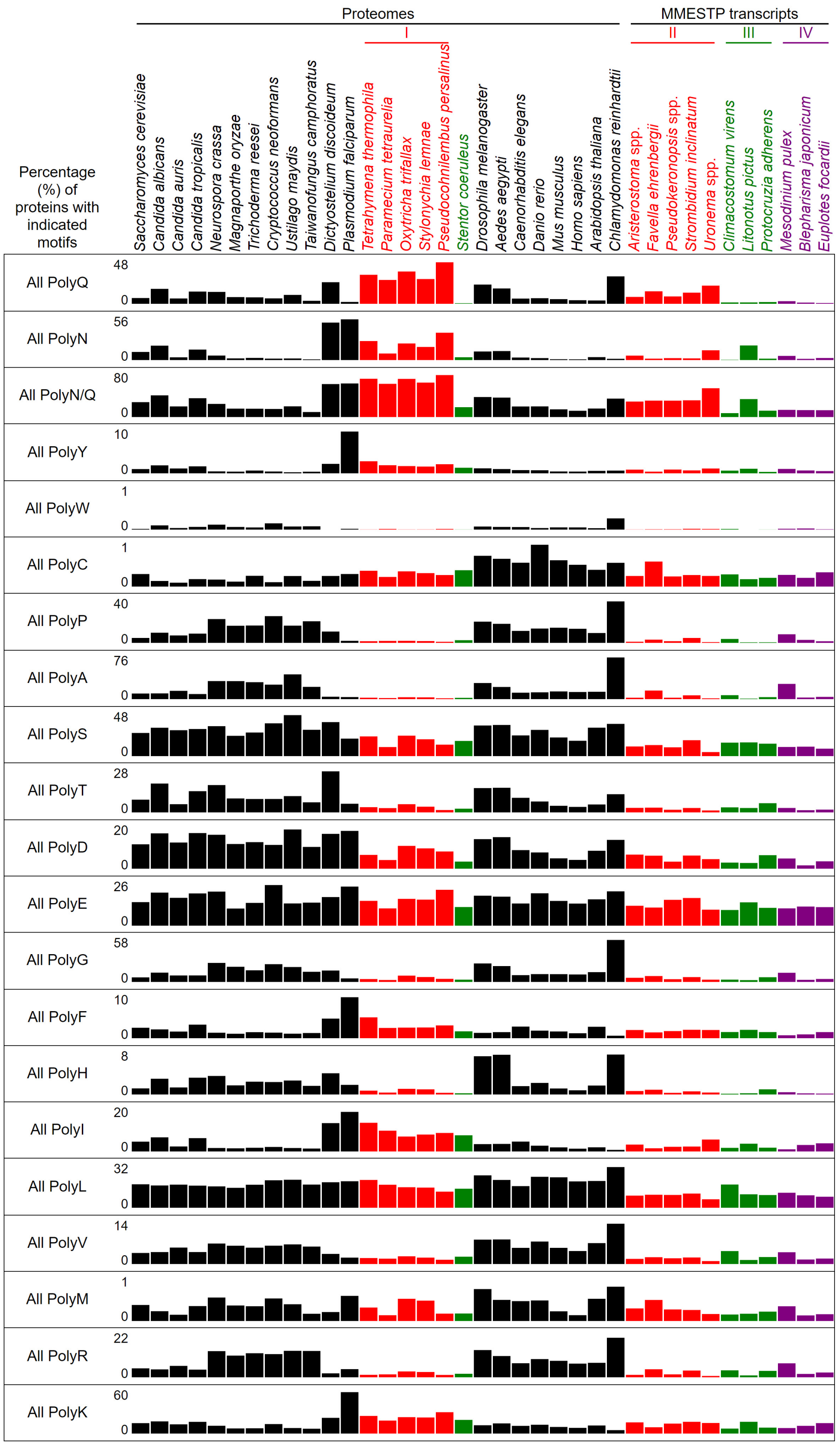


**Figure S3. Percentages of proteins with indicated polyX motifs in 37 different eukaryotes.**

Table S1. Plasmids used in this study

| Plasmid number | Description |
|---|---|
| pTFW2592 | pYC2/NT-C, The mock control. |
| pTFW9957 | $P_{RAD51}$*-LacZ-NVH* |
| pTFW9958 | $P_{RAD51}$*-RAD51-NTD(1-66 a.a.)- LacZ-NVH* |
| pTFW9962 | $P_{RAD51}$*-RAD53-SCD1(1-29 a.a.)-LacZ-NVH* |
| pTFW9963 | $P_{RAD51}$*-HOP1-SCD(258-324 a.a.)-LacZ-NVH* |
| pTFW10027 | $P_{RAD51}$*-SML1-NTD(1-27 a.a.)-LacZ-NVH* |
| pTFW10028 | $P_{RAD51}$*-SML1-NTD(1-50 a.a.)-LacZ-NVH* |
| pTFW10005 | $P_{RAD51}$*-SUP35-PND(1-39 a.a.)-LacZ-NVH* |
| pTFW10221 | $P_{RAD51}$*-SUP35-PFD(1-114 a.a.)-LacZ-NVH* |
| pTFW10169 | $P_{RAD51}$*-NEW1-NPD(1-156 a.a.)-LacZ-NVH* |
| pTFW10177 | $P_{RAD51}$*-URE2-UPD(1-91 a.a.)-LacZ-NVH* |
| pTFW9959 | $P_{RAD51}$*-rad51-NTD-3SA-LacZ-NVH* |
| pTFW10095 | $P_{RAD51}$*-rad51-NTD-8STA-LacZ-NVH* |
| pTFW10096 | $P_{RAD51}$*-rad51-NTD-11STA-LacZ-NVH* |
| pTFW9993 | $P_{RAD51}$*-rad51-NTD-6SQA-LacZ-NVH* |
| pTFW9994 | $P_{RAD51}$*-rad51-NTD-9SQA-LacZ-NVH* |
| pTFW9995 | $P_{RAD51}$*-rad51-NTD-12SQA-LacZ-NVH* |
| pTFW10010 | $P_{RAD51}$*-rad51-NTD-3QA -LacZ-NVH* |
| pTFW10254 | $P_{RAD51}$*-rad51-NTD-9QA-LacZ-NVH* |
| pTFW10255 | $P_{RAD51}$*-rad51-NTD-4NA-LacZ-NVH* |
| pTFW10256 | $P_{RAD51}$*-rad51-NTD-13QNA-LacZ-NVH* |
| pTFW10000 | $P_{RAD51}$*-rad53-SCD1-5STA-LacZ-NVH* |
| pTFW10001 | $P_{RAD51}$*-rad53-SCD1-7QA-LacZ-NVH* |
| pTFW10002 | $P_{RAD51}$*-rad53-SCD1-12STQA-LacZ-NVH* |
| pTFW10034 | $P_{RAD51}$*-sup35-PND-1SA-LacZ-NVH* |
| pTFW10101 | $P_{RAD51}$*-sup35-PND-3SA-LacZ-NVH* |
| pTFW10081 | $P_{RAD51}$*-sup35-PND-3QA-LacZ-NVH* |
| pTFW10080 | $P_{RAD51}$*-sup35-PND-5QA-LacZ-NVH* |
| pTFW10082 | $P_{RAD51}$*-sup35-PND-8QA-LacZ-NVH* |
| pTFW10141 | $P_{RAD51}$*-sup35-PND-15SQA-LacZ-NVH* |
| pTFW10142 | $P_{RAD51}$*-sup35-PND-9NA-LacZ-NVH* |
| pTFW10143 | $P_{RAD51}$*-sup35-PND-24SQNA-LacZ-NVH* |
| pTFW9974 | $P_{RAD51}$*-GFP-NVH* |
| pTFW9975 | $P_{RAD51}$*-RAD53-SCD1-GFP-NVH* |
| pTFW9976 | $P_{RAD51}$*-GST-NVH* |
| pTFW9977 | $P_{RAD51}$*-RAD53-SCD1-GST-NVH* |
| pTFW10003 | $P_{RAD51}$*- GSTnd-NVH* |
| pTFW10004 | $P_{RAD51}$*-RAD53-SCD1-GSTnd-NVH* |

Table S2. *S. cerevisiae* strains used in this study

| Name | Genotype |
|---|---|
| WHY13008 | *MATa, ho, leu2, ura3, his4-X::LEU2-(NgoMIV;+ori)-URA3, ERG1(SpeI), RAD51::hphMX4* |
| WHY13283 | *MATa, ho, leu2, ura3, his4-X::LEU2-(NgoMIV;+ori)-URA3, ERG1(SpeI), rad51Δ::hphMX4* |
| WHY13416 | *MATa, ho, leu2, ura3, his4-X::LEU2-(NgoMIV;+ori)-URA3, ERG1(SpeI), rad51ΔN::hphMX4* |
| WHY13744 | *MATα, ho::LYS2, leu2, ura3, lys2, HIS4::LEU2-(BamHI;+ori), Erg1(SalI), dmc1::kanMX4, RAD53$^{SCD1}$-rad51ΔN::hphMX4* |
| WHY13743 | *MATa, ho, leu2, ura3, his4-X::LEU2-(NgoMIV;+ori)-URA3, ERG1(SpeI), dmc1::kanMX4, rad53$^{SCD1-5STA}$-rad51ΔN::hphMX4* |
| WHY13741 | *MATα, ho::LYS2, leu2, ura3, lys2, HIS4::LEU2-(BamHI;+ori), ERG1(SalI), SUP35$^{PND}$-rad51ΔN::hphMX4* |
| WHY10271 | *MATa, ho::hisG, lys2, leu2::hisG, arg4-nsp, ura3* |
| WHY13970 | *MATa his3Δ1, leu2Δ0, met15Δ0, ura3Δ0* |
| WHY13785 | *MATa his3Δ1, leu2Δ0, met15Δ0, ura3Δ0, hsp104::kanMX4* |
| WHY14126 | *MATa his3Δ1, leu2Δ0, met15Δ0, ura3Δ0, new1::kanMX4* |
| WHY14129 | *MATa his3Δ1, leu2Δ0, met15Δ0, ura3Δ0, doa4::kanMX4* |
| WHY14227 | *MATa his3Δ1, leu2Δ0, met15Δ0, ura3Δ0, doa1::kanMX4* |
| WHY13989 | *MATa his3Δ1, leu2Δ0, met15Δ0, ura3Δ0, san1::kanMX4* |
| WHY14132 | *MATa his3Δ1, leu2Δ0, met15Δ0, ura3Δ0, oaz1::kanMX4* |

Table S3. The oligonucleotide primers used for g-qPCR and RT-qPCR

| Gene | Primer Polarity | Name | Sequence |
|---|---|---|---|
| *ACT1* | Sense | PA11240 | 5’-CCACCACTGCTGAAAGAGAAATTGT-3’ |
| | Antisense | PA11241 | 5’-CTTGACCATCTGGAAGTTCGTAGGA-3’ |
| *LacZ* | Sense | PA11246 | 5’-CCGCCGTTTGTTCCCACGGA-3’ |
| | Antisense | PA11247 | 5’-CCATCACCGCGAGGCGGTTT-3’ |

Table S4. The JavaScript software programs used in this study are available at Github (https://github.com/tfwangasimb/AS-Q-rich-motif)

| Program name | Purpose |
|---|---|
| AS-aa-content | Determination of proteome-wide contents of 20 different amino acids |
| AS-codon-usage | Determination of proteome-wide frequency of 64 genetic codons |
| AS-Finder-SCD | Proteome-wide search of the SCD motifs |
| AS-Finder-7polyX | Proteome-wide search of the 7 different types of polyX motifs |
| AS-Xcontent-7polyX | Determination of the ratios of the overall number of X residues for each of the seven polyX motifs relative to those in the entire proteome of each species |
| AS-GOfuncR-FWER | Statistical analysis of GO enrichment datasets |
